# Cortical synapses of the world’s smallest mammal: an FIB/SEM study in the Etruscan shrew

**DOI:** 10.1101/2022.06.06.494946

**Authors:** Lidia Alonso-Nanclares, J. Rodrigo Rodríguez, Ángel Merchan-Perez, Juncal González-Soriano, Sergio Plaza-Alonso, Nicolás Cano-Astorga, Robert K. Naumann, Michael Brecht, Javier DeFelipe

**Author notes:** **Corresponding author:** Javier DeFelipe.

## Abstract

The main aim of the present study was to determine if synapses from the exceptionally small brain of the Etruscan shrew show any peculiarities compared to the much larger human brain. This study constitutes the first description of the Etruscan shrew synaptic characteristics using Focused Ion Beam/Scanning Electron Microscopy (FIB/SEM). We analyzed the synaptic density and a variety of structural characteristics of 7,239 3D reconstructed synapses, obtaining the following major results: (i) cortical synaptic density was very high, particularly in layer I; (ii) the vast majority of synapses were excitatory, with the highest proportion found in layer I; (iii) excitatory synapses were larger than inhibitory synapses in all layers except in layer VI; and (iv) synapses were either randomly distributed in space or showed a slight tendency to be organised in a regular arrangement. Some of these general synaptic characteristics are remarkably similar to those found in the human cerebral cortex. However, the cortical volume of the human brain is about 50,000 times larger than the cortical volume of the Etruscan shrew, while the total number of cortical synapses in human is *only* 20,000 times the number of synapses in the shrew, and synaptic junctions are 35% smaller in the Etruscan shrew. Thus, the differences in the number and size of synapses cannot be attributed to a brain size scaling effect but rather to adaptations of synaptic circuits to particular functions. The present work provides a quantitative dataset from the Etruscan shrew synapses — not only contributing to the knowledge of the ultrastructure of the mammalian cortex, but also identifying common and differing principles of synaptic organization.

## INTRODUCTION

The Etruscan shrew (*Suncus etruscus*; also known as the Etruscan pygmy shrew or the white-toothed pygmy shrew) is the smallest known terrestrial mammal by mass, weighing only about 1.8 g on average (Fons et al., 1984; Jürgens, 2002). This tiny mammal has a body length of about 4 cm excluding the tail, and its brain is the smallest of all mammalian species, with a brain mass of only about 0.06 g (e.g., Fons et al., 1984). Furthermore, the neocortex of the Etruscan shrew is the thinnest among all mammals, with a thickness of only 400–500 μm (Stolzenburg et al., 1989; Roth-Alpermann et al., 2010; Naumann et al., 2012) and an extremely high density of neurons — as high as 170,000 neurons per mm^3^ (Stolzenburg et al., 1989). Another peculiarity of these animals is their very fast metabolism — they have been reported to eat up to 6 times their own body weight per day (Brecht et al., 2011). The Etruscan shrew can hunt animals the same size as itself, showing remarkable speed and accuracy to recognize prey shape based on whisker-mediated tactile cues (Brecht et al., 2011; Naumann et al., 2012).

The neocortex of the Etruscan shrew is a cytoarchitectonically heterogeneous sheet with distinct cortical areas. Around 13 cortical areas have been distinguished (Naumann et al., 2012), which is a large number considering that the human cortex is 50,000 times larger (Ribeiro et al., 2013) and has about 150–200 distinct areas (Amunts & Zilles, 2015).

Sensory cortical areas in the Etruscan shrew occupy a large portion of the total cortical volume (Brecht et al., 2011). In fact, 25% of the neocortical neurons are located in the somatosensory cortex (Naumann et al., 2012), pointing to the key functional importance of the somatosensory cortex. Around 75% of the shrew cortex responds to tactile stimuli (Roth-Alpermann et al., 2010), which mostly relies on somatosensory cortical regions. As mentioned above, the Etruscan shrew has a highly specialized system of tactile object recognition based on its whiskers, which is critical for prey capture, and, consequently, for survival (Anjum et al., 2006; Roth-Alpermann et al., 2010).

The aim of the present study was to analyze the primary somatosensory cortex of the Etruscan shrew at the ultrastructural level, to determine whether the cortical synapses show any peculiarities that may be related to its small brain size, thin cortex and high neuronal density. For this purpose, we examined all cortical layers (I, II, III, IV, V, VI) using Focused Ion Beam/Scanning Electron Microscopy (FIB/SEM) to obtain quantitative information on cortical synapses. Specifically, we analyzed the synaptic density of 7,239 3D-reconstructed synapses as well as a variety of their structural characteristics including: the type of synapse (asymmetric or symmetric, corresponding to excitatory and inhibitory synapses, respectively), the size of each 3D reconstructed synapse, as well as the 3D spatial distribution of each synapse. The results are discussed comparing with data obtained from the human cerebral cortex using the same technology (Domínguez-Álvaro et al., 2018; 2021a; Cano-Astorga et al., 2021). From an evolutionary point of view, it is of particular interest to compare the cortical synaptic organization of this extremely small mammal with that of the much larger human brain (Hofman, 1988), whose synaptic organization is thought to have reached the highest level of complexity.

## MATERIALS AND METHODS

### Tissue preparation

Brain tissue from 3 male (8-, 12- and 20-month-old) Etruscan shrews (*Suncus etruscus*) were used for this study. The animals were briefly anesthetized using isoflurane and subsequently given an intraperitoneal injection of 20% urethane prior to intracardial perfusion of a fixative solution containing 2% paraformaldehyde and 2.5% glutaraldehyde in 0.1 M phosphate buffer. The brain was extracted from the skull and fixed overnight in the same fixative solution at 4°C. Brain sections (150 μm thick) were obtained coronally (Vibratome Sectioning System, VT1200S Vibratome, Leica Biosystems, Germany), and processed following the protocols described below. All animals were handled in accordance with the guidelines for animal research set out in European Community Directive 2010/63/EU.

### Electron microscopy

Brain sections were post-fixed for 24h in a solution containing 2% paraformaldehyde, 2.5% glutaraldehyde (TAAB, G002, UK) and 0.003% CaCl_2_ (Sigma, C-2661-500G, Germany) in sodium cacodylate (Sigma, C0250-500G, Germany) buffer (0.1M). The sections were treated with 1% OsO_4_ (Sigma, O5500, Germany) and 0.003% CaCl_2_ in sodium cacodylate buffer (0.1M) for 1h at room temperature. They were then stained with 1% uranyl acetate (EMS, 8473, USA), dehydrated and flat-embedded in Araldite (TAAB, E021, UK) for 48h at 60°C (DeFelipe & Fairén, 1993). The embedded sections were then glued onto a blank Araldite block. Semithin sections (2 μm thick) were obtained from the blocks and stained with 1% toluidine blue (Merck, 115930, Germany) in 1% sodium borate (Panreac, 141644, Spain). For each block, the last semithin section (corresponding to the section immediately adjacent to the block surface) was examined under light microscope and photographed to accurately locate the neuropil regions to be examined by electron microscopy (Fig. 1).

**Figure 1.**
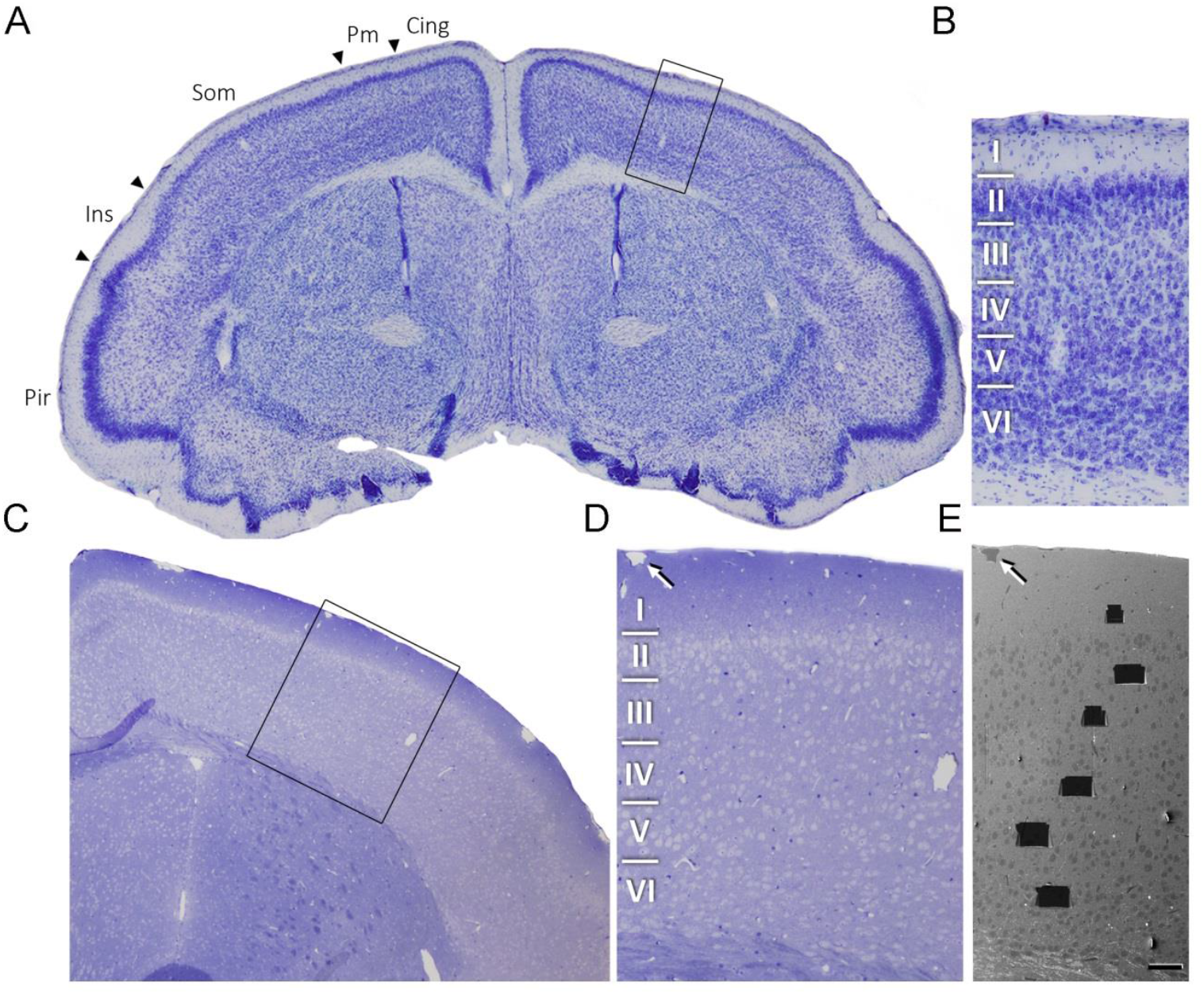
Correlative light-electron microscopy study of the Etruscan shrew cerebral cortex. **(A)** Low power photograph of a 150 μm Nissl stained coronal vibratome section of the Etruscan shrew brain. The delimitation of cortical areas and layers is based on Naumann et al. (2012). **(B)** Higher magnification of the boxed area in A, showing the laminar pattern of Som cortex (layers I to VI are indicated). **(C)** 1 μm-thick semithin section stained with toluidine blue. **(D)** Higher magnification of the boxed area in C, showing delimitated layers based on the staining pattern. The semithin section is adjacent to the block for FIB/SEM imaging. **(E)** SEM image illustrating the block surface with trenches made in the neuropil (one per layer). Arrows in D and E point to the same blood vessel, showing that the exact location of the region of interest was accurately determined. Scale bar shown in E represents 200 μm in A, 60 μm in B, 240 μm in C, 100 μm in D and 55 μm in E. Cing – Cingulate Cortex; Pm – Parietal Medial Cortex; Som – Somatosensory Cortex; Ins – Insular Cortex; Pir – Piriform Cortex).

### Three-dimensional electron microscopy

Images were obtained from the neuropil, which is where the vast majority of synapses are found (DeFelipe et al., 1999). The neuropil is composed of axons, dendrites and glial processes, so the samples did not contain cell somata, proximal dendrites in the immediate vicinity of the soma, or blood vessels.

Three-dimensional brain tissue samples of the somatosensory cortex were obtained using a Neon40 EsB electron microscope (Carl Zeiss NTS GmbH, Oberkochen, Germany). This instrument combines a high-resolution field emission SEM column with a focused gallium ion beam (FIB), which mills the sample surface, removing thin layers of material on a nanometer scale. After removing each slice (20 nm thick), the milling process was paused, and the freshly exposed surface was imaged with a 1.7 kV acceleration potential using the in-column energy selective backscattered (EsB) electron detector. The milling and imaging processes were sequentially repeated, and long series of images were acquired through a fully automated procedure, thus obtaining a stack of images that represented a three-dimensional sample of the tissue (Merchan-Perez et al., 2009). Eighteen samples (stacks of images) of the neuropil from the somatosensory cortex were obtained in the six layers (one sample per layer and per animal, in layers I, II, III, IV, V and VI).

Image resolution in the xy plane was 4.652nm/pixel. Resolution in the z axis (section thickness) was 20 nm and image sizes were 2048 × 1536 pixels. These parameters allowed a field of view where synaptic junctions could be clearly identified, within a reasonable image acquisition timeframe (approximately 12 h per stack of images). The number of sections per stack ranged from 200 to 301 (accumulative total: 4,335 sections). The volumes of the stacks ranged from 339 to 527 μm3, and a total volume of 7,460 μm3 was sampled (considering the corrected volume that accounted for tissue shrinkage). All measurements were corrected for the tissue shrinkage that occurs during the processing of sections (Merchan-Perez et al., 2009). To estimate the shrinkage in our samples, we photographed and measured the area of the brain sections with ImageJ (ImageJ 1.51; NIH, USA), both before and after processing for electron microscopy. The section area values after processing were divided by the values before processing to obtain the volume, area, and linear shrinkage factors (Oorschot et al., 1991), yielding correction factors of 0.803, 0.864 and 0.929, respectively. Nevertheless, in order to compare with previous studies —in which no correction factors had been included or such factors were estimated using other methods— in the present study, we provide both sets of data.

### Three-dimensional analysis of synapses

Stacks of images obtained by the FIB/SEM were analyzed using EspINA software (EspINA Interactive Neuron Analyzer, 2.1.9; https://cajalbbp.es/espina/). As previously discussed (Merchan-Perez et al., 2009), there is a consensus for classifying cortical synapses into asymmetric synapses (AS; or type I) and symmetric synapses (SS; or type II). The main characteristic distinguishing these synapses is the prominent or thin post-synaptic density, respectively (Gray, 1959; Colonnier, 1968; Peters et al., 1991; Fig. 2;

**Figure 2.**
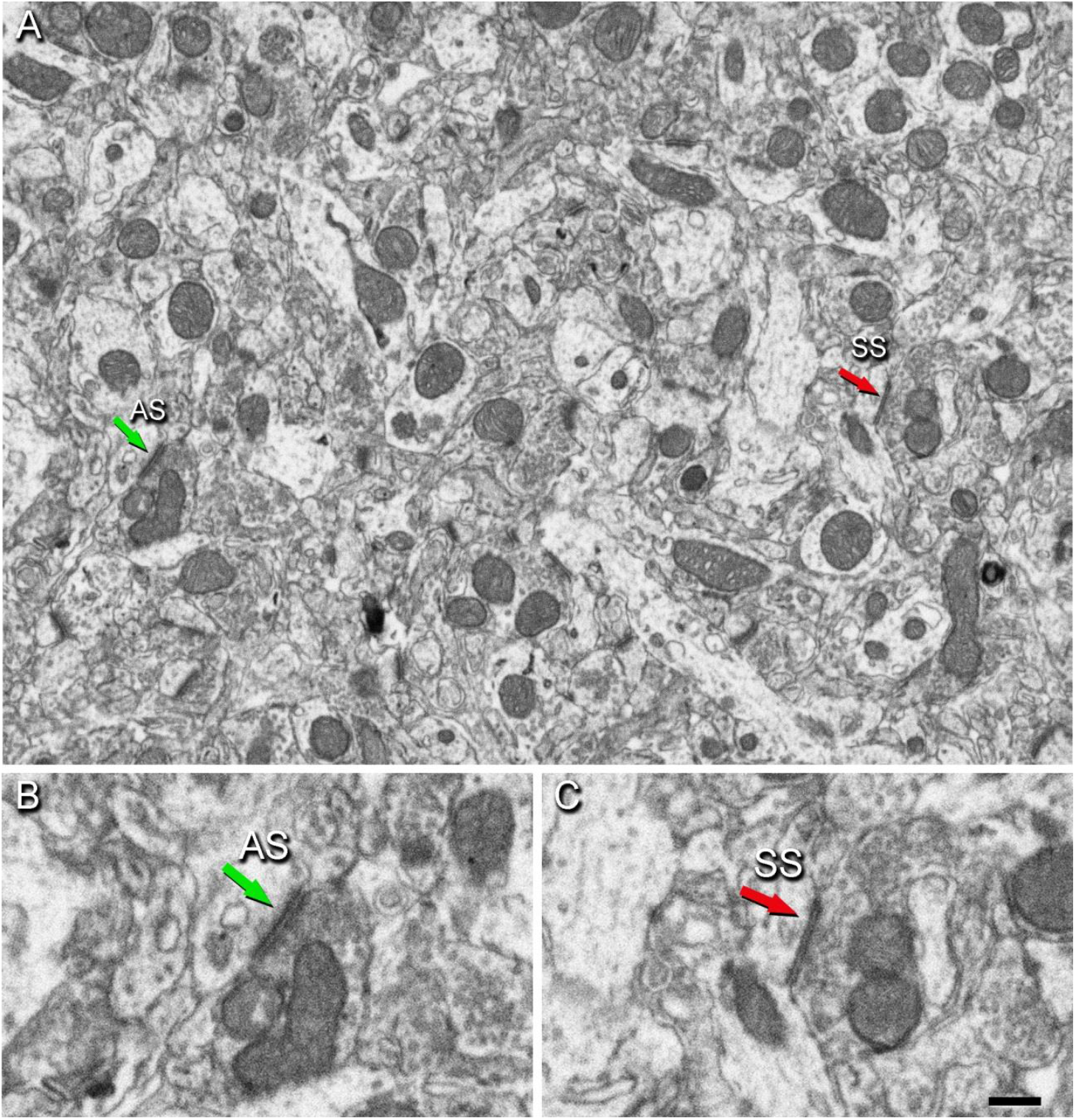
Images of neuropil in layer III of Etruscan shrew somatosensory cortex obtained by FIB/SEM. **(A)** Two synapses are indicated as examples of asymmetric (AS, green arrow) and symmetric (SS, red arrow) synapses. **(B, C)** Higher magnification of AS (B) and SS (C) indicated in (A). Synapse classification was based on the examination of the full sequence of serial images (Supplementary Fig. 1). Scale bar in C represents 500 nm in A, and 250 nm in B and C.

Supplementary Fig. 1). Also, these two types of synapses are associated with different functions: AS are mostly glutamatergic and excitatory, while SS are mostly GABAergic and inhibitory (DeFelipe & Fariñas, 1992; Houser et al., 1984; Ascoli et al., 2008).

Nevertheless, in single sections, the synaptic cleft and the pre- and post-synaptic densities are often blurred if the plane of the section does not pass at right angles to the synaptic junction. Since the software EspINA allows navigation through the stack of images, it was possible to unambiguously identify every synapse as AS or SS, based on the thickness of the postsynaptic density (PSD) (Merchan-Perez et al., 2009).

EspINA provided the 3D reconstruction of every synapse and allowed the application of an unbiased 3D counting frame (CF), which is a rectangular prism enclosed by three acceptance planes and three exclusion planes marking its boundaries. All synapses within the CF were counted, as were those intersecting any of the acceptance planes, while synapses that were outside the CF, or intersecting any of the exclusion planes, were not counted (Fig. 3). Thus, the number of synapses per unit volume was calculated directly by dividing the total number of synapses counted by the volume of the CF (Merchan-Pérez et al., 2009), in all 18 stacks of images.

**Figure 3.**
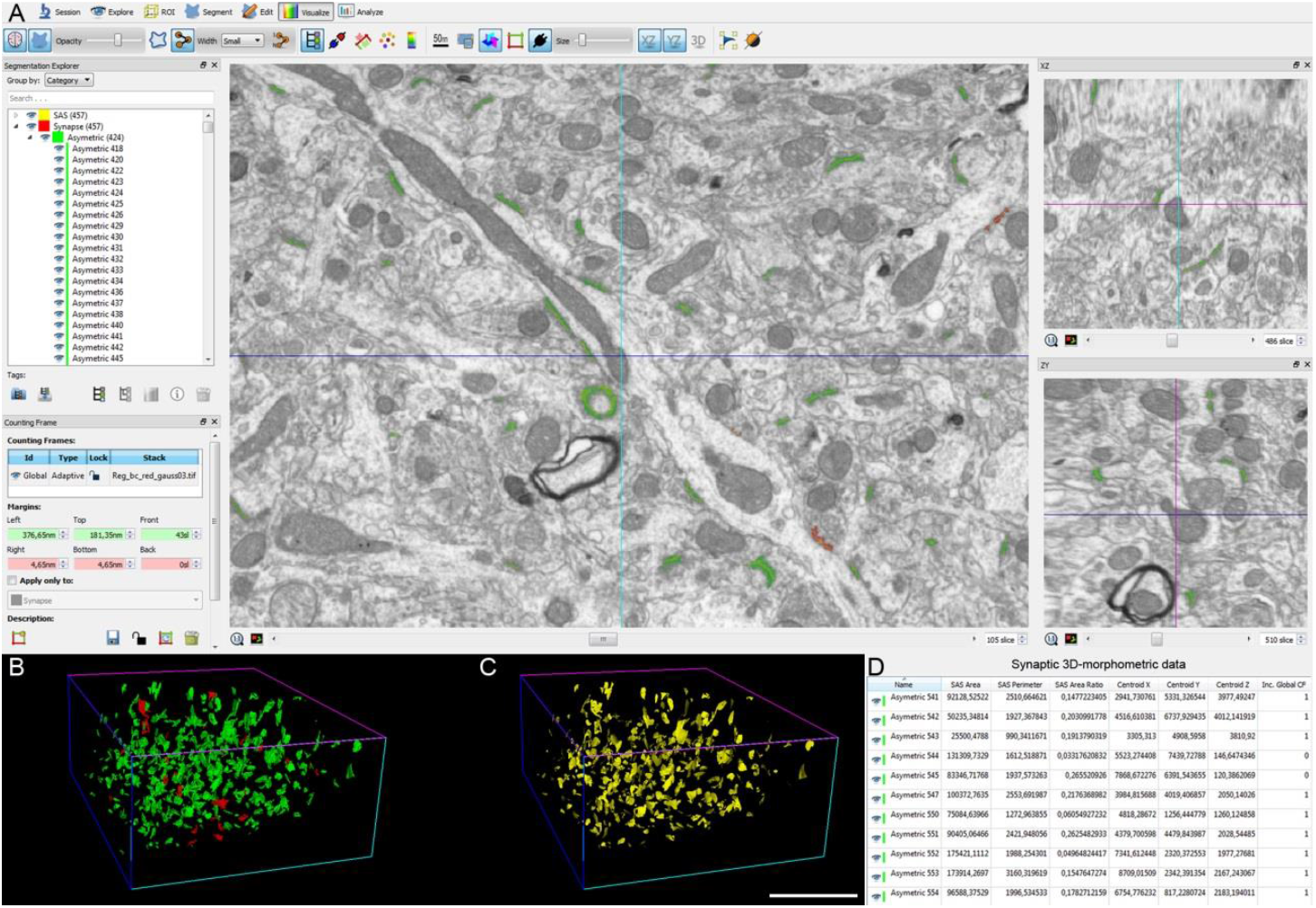
Screenshot of the EspINA software user interface. **(A)** In the main window, the sections are viewed through the xy plane (as obtained by FIB/SEM microscopy). The other two orthogonal planes, yz and xz, are also shown in adjacent windows (on the right). **(B)** 3D reconstructions of segmented AS (green) and SS (red). **(C)** Computed SAS for each reconstructed synapse (yellow). **(D)** Table of synaptic 3D morphometric data from AS automatically obtained by EspINA software. Scale bar in C represents 5 μm in B and C.

Synaptic size was calculated using the Synaptic Apposition Surface (SAS), which was automatically extracted by EspINA (Fig. 3). The SAS represents both the active zone (presynaptic density) and the PSD, resulting in a functionally relevant measurement of the synaptic size (Morales et al., 2013). Estimations of the SAS were made for each individually 3D reconstructed complete synapse in all FIB/SEM stacks, with the SAS area providing a reliable synaptic size measurement.

### Spatial distribution analysis of synapses

In addition, the positions of the centers of gravity (centroids) of each reconstructed synapse were also calculated by EspINA in all FIB/SEM stacks of images.

To analyze the spatial distribution of synapses, Spatial Point Pattern analysis was performed on the centroids as described elsewhere (Antón-Sánchez et al., 2014; Merchan-Perez et al., 2014). Briefly, we compared the actual position of synapse centroids with the Complete Spatial Randomness (CSR) model — a random spatial distribution model which defines a situation where a point is equally likely to occur at any location within a given volume. To do this, we generated an envelope simulating 99 instances of random distributions of the same number of points as our experimental sample.

Then, for each of the 18 FIB/SEM stacks of images, we calculated three functions commonly used for spatial point pattern analysis: F, G and K functions. When these functions lay within the envelope, we concluded that the distributions of synapses were random. Otherwise, the distribution of points may be clustered (when points are closer to each other than expected by chance) or regular (when points tend to separate from each other further that expected by chance). The F function, also known as the empty space function or the point-to-event distribution, is the cumulative distribution of distances between the centroids of synapses and the closest point in a regularly spaced grid of points superimposed over the sample. The G function, also called the nearest-neighbor distance cumulative distribution function or the event-to-event distribution, is the cumulative distribution of distances between each centroid and its nearest neighbor. The K function is also called the reduced second moment function or Ripley’s function. An estimation of the K function is given by the mean number of points within a sphere of increasing radius centered on each sample centroid. See Merchan-Perez et al. (2014) and Anton-Sanchez et al. (2014) for examples of studies in which this methodology was used to investigate the spatial distribution of synapses. The present study was carried out using the Spatstat package and R Project program (Baddeley et al., 2015).

### Statistical analysis

To study whether there were significant differences between synaptic characteristics among the different layers, we performed a multiple mean comparison test on the 18 samples of the six cortical layers. If the necessary assumptions for ANOVA were not satisfied (the normality and homoscedasticity criteria were not met), we used the Kruskal–Wallis test (KW) and the Mann–Whitney test (MW) for pair-wise comparisons. χ^2^ tests were used for contingency table analysis. Frequency distribution analysis of the SAS area was performed using Kolmogorov-Smirnov (KS) nonparametric test. Statistical studies were performed with the GraphPad Prism statistical package (Prism 9.00 for Windows, GraphPad Software Inc., USA), Spatstat package for R Project program (Baddeley et al., 2015) and Easyfit Professional 5.5 (MathWave Technologies).

## RESULTS

The following results were obtained in the neuropil, so they represent synapses located among cell bodies, excluding perisomatic synapses and synapses established on thick proximal dendritic trunks.

### Synaptic Density

The number of synapses per volume was calculated in the 18 stacks of images obtained from 3 animals, in 6 layers per animal. A total of 9,033 synapses were individually identified and reconstructed in 3D. Of these, 7,239 synapses were analyzed after discarding synapses that were truncated by the margins of the stack or those touching the exclusion edges of the counting frame (CF). Summing all the CFs that were applied yielded a total volume of 5,578μm^3^ (Table 1). The synaptic density values were obtained by dividing the total number of synapses included within each CF by its total volume. Since the synapses were fully reconstructed in 3D, it was possible to classify them as AS and SS based on the thickness of their postsynaptic densities (PSDs), allowing us to compute the densities and proportions of AS and SS in each cortical layer (Merchan-Perez et al., 2009).

**Table 1.** Accumulated synaptic data per animal. Data in parentheses are not corrected for shrinkage. AS: asymmetric synapses; SAS: synaptic apposition surface; SEM: standard error of the mean; SD: standard deviation; SS: symmetric synapses.

| Animal | No. of AS | No. of SS | Total no. of synapses | % AS (mean±SD) | % SS (mean±SD) | Total Analyzed Volume (μm<sup>3</sup>) | No. of AS/μm<sup>3</sup> (mean ± SD) | No. of SS/μm<sup>3</sup> (mean ± SD) | Total no. of synapses/μm<sup>3</sup> (mean ± SD) | Area of SAS AS (nm<sup>2</sup>; mean ± SEM) | Area of SAS SS (nm<sup>2</sup>; mean ± SEM) | Intersynaptic distance (nm; mean± SD) |
| --- | --- | --- | --- | --- | --- | --- | --- | --- | --- | --- | --- | --- |
| MS1 | 1750 | 249 | 1999 | 87.4±5.3 | 12.6±5.3 | 1831 (1469) | 0.95±0.22 (1.19±0.27) | 0.13±0.05 (0.16±0.06) | 1.08±0.21 (1.35±0.26) | 77249±5557 (66699±4798) | 63774±3950 (55064±3411) | 629±51 (584±47) |
| MS2 | 2223 | 209 | 2432 | 91.4±3.1 | 8.6±3.1 | 1957 (1570) | 1.2±0.4 (1.49±0.5) | 0.10±0.03 (0.13±0.04) | 1.30±0.39 (1.62±0.49) | 70672±4102 (61020±3542) | 53290±2813 (46012±2429) | 574±54 (533±50) |
| MS3 | 2591 | 217 | 2808 | 92.1±1.5 | 7.9±1.5 | 1790 (1436) | 1.43±0.29 (1.78±0.37) | 0.12±0.03 (0.15±0.04) | 1.55±0.31 (1.93±0.39) | 74067±4380 (63952±3782) | 64071±5507 (55321±4755) | 570±51 (529±47) |
| Total | 6564 | 675 | 7239 | 90.3±2.5 | 9.7±2.5 | 5578 (4475) | 1.19±0.24 (1.49±0.29) | 0.12±0.01 (0.15±0.02) | 1.31±0.23 (1.63±0.29) | 73996±3289 (63890±2840) | 60378±6140 (52133±5302) | 591±33 (549±31) |

The overall synaptic density —obtained by averaging all layers and animals— was 1.31 synapses/μm^3^ (Table 1). The total synaptic density and AS density reached the highest values in layer I (1.70 and 1.62 synapses/μm^3^, respectively), and the lowest values in layer VI (1.01 and 0.91 synapses/μm^3^, respectively; Fig. 4A, Table 2, Supplementary Table 1). Regarding SS, the density was highest in layer III and lowest in layer I (Table 2).

**Figure 4.**
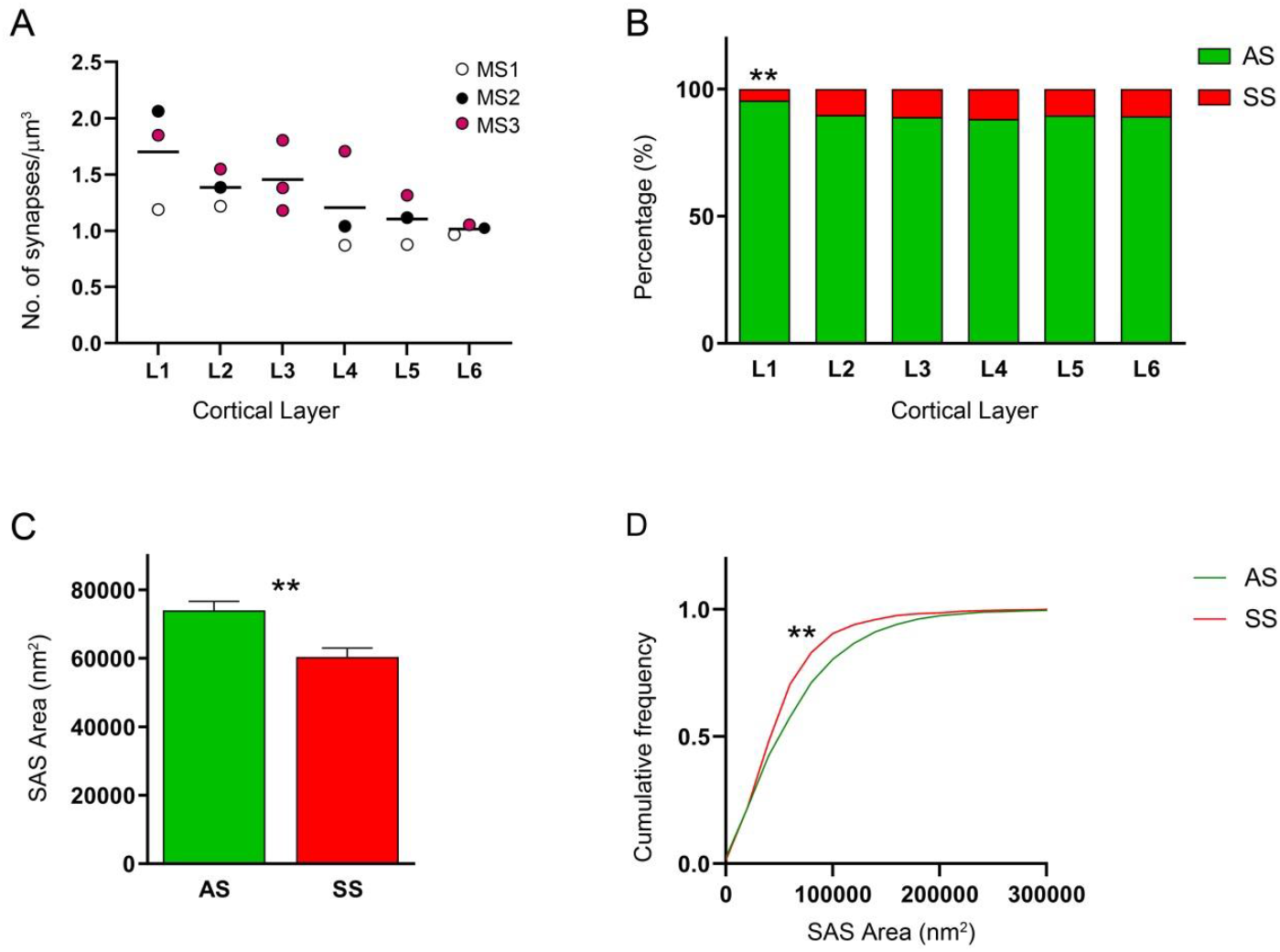
Plots of the synaptic analysis of the Etruscan shrew somatosensory cortex. **(A)** Mean of the overall synaptic density from each layer. Different colors correspond to each analyzed animal, as denoted in the upper right-hand corner. **(B)** Proportion of AS and SS per layer expressed as percentages, showing that layer 1 was different from the other layers (χ^2^; P <0.0001). **(C)** Mean SAS area per synaptic type shows larger synaptic size of AS compared to SS (MW, p=0.0015). **(D)** Cumulative frequency distribution graph of SAS area illustrating that small SS (red) were more frequent (KS, p<0.0001) than small AS (green). Asterisks indicate statistically significant differences.

**Table 2.**
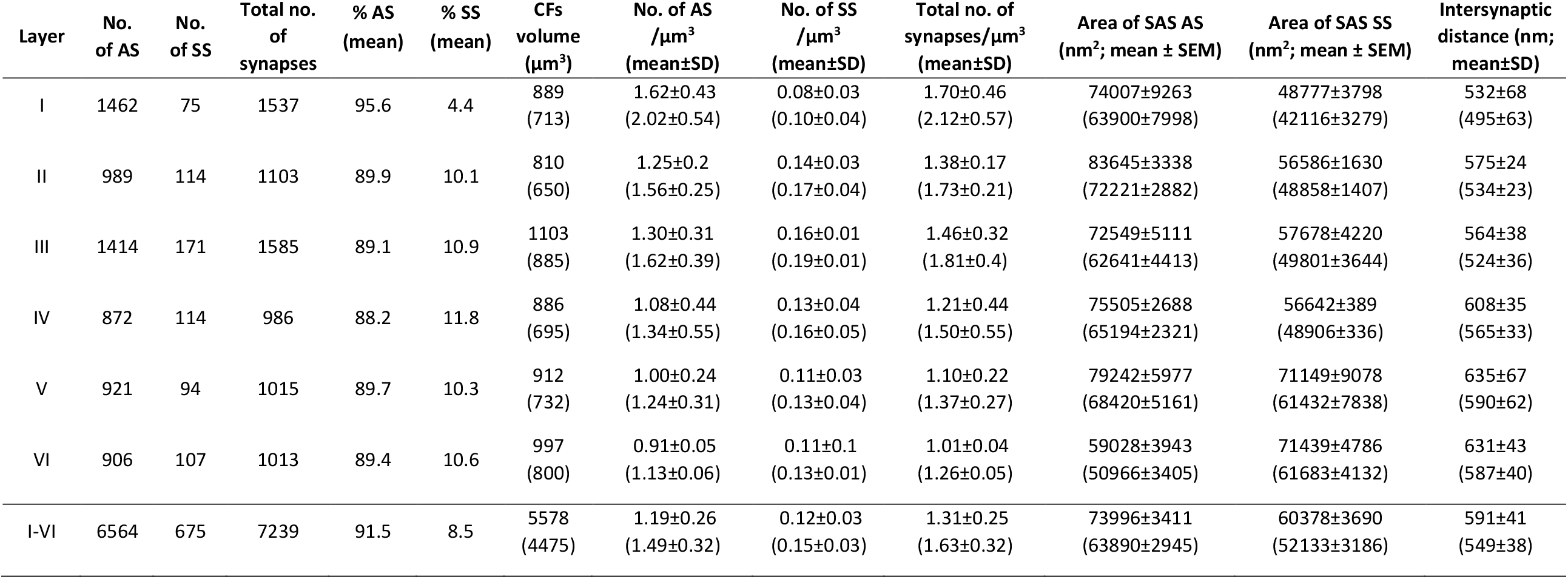
Synaptic data per layer. Data in parentheses are not corrected for shrinkage. AS: asymmetric synapses; CF: counting frame; SAS: synaptic apposition surface; SEM: standard error of the mean; SD: standard deviation; SS: symmetric synapses.

| Layer | No. of AS | No. of SS | Total no. of synapses | % AS (mean) | % SS (mean) | CFs volume ($\mu\text{m}^3$) | No. of AS /$\mu\text{m}^3$ (mean±SD) | No. of SS /$\mu\text{m}^3$ (mean±SD) | Total no. of synapses/$\mu\text{m}^3$ (mean±SD) | Area of SAS AS ($\text{nm}^2$; mean ± SEM) | Area of SAS SS ($\text{nm}^2$; mean ± SEM) | Intersynaptic distance (nm; mean±SD) |
| --- | --- | --- | --- | --- | --- | --- | --- | --- | --- | --- | --- | --- |
| I | 1462 | 75 | 1537 | 95.6 | 4.4 | 889 (713) | 1.62±0.43 (2.02±0.54) | 0.08±0.03 (0.10±0.04) | 1.70±0.46 (2.12±0.57) | 74007±9263 (63900±7998) | 48777±3798 (42116±3279) | 532±68 (495±63) |
| II | 989 | 114 | 1103 | 89.9 | 10.1 | 810 (650) | 1.25±0.2 (1.56±0.25) | 0.14±0.03 (0.17±0.04) | 1.38±0.17 (1.73±0.21) | 83645±3338 (72221±2882) | 56586±1630 (48858±1407) | 575±24 (534±23) |
| III | 1414 | 171 | 1585 | 89.1 | 10.9 | 1103 (885) | 1.30±0.31 (1.62±0.39) | 0.16±0.01 (0.19±0.01) | 1.46±0.32 (1.81±0.4) | 72549±5111 (62641±4413) | 57678±4220 (49801±3644) | 564±38 (524±36) |
| IV | 872 | 114 | 986 | 88.2 | 11.8 | 886 (695) | 1.08±0.44 (1.34±0.55) | 0.13±0.04 (0.16±0.05) | 1.21±0.44 (1.50±0.55) | 75505±2688 (65194±2321) | 56642±389 (48906±336) | 608±35 (565±33) |
| V | 921 | 94 | 1015 | 89.7 | 10.3 | 912 (732) | 1.00±0.24 (1.24±0.31) | 0.11±0.03 (0.13±0.04) | 1.10±0.22 (1.37±0.27) | 79242±5977 (68420±5161) | 71149±9078 (61432±7838) | 635±67 (590±62) |
| VI | 906 | 107 | 1013 | 89.4 | 10.6 | 997 (800) | 0.91±0.05 (1.13±0.06) | 0.11±0.1 (0.13±0.01) | 1.01±0.04 (1.26±0.05) | 59028±3943 (50966±3405) | 71439±4786 (61683±4132) | 631±43 (587±40) |
| I-VI | 6564 | 675 | 7239 | 91.5 | 8.5 | 5578 (4475) | 1.19±0.26 (1.49±0.32) | 0.12±0.03 (0.15±0.03) | 1.31±0.25 (1.63±0.32) | 73996±3411 (63890±2945) | 60378±3690 (52133±3186) | 591±41 (549±38) |

The general proportion of AS:SS, computed for all animals and layers collected was approximately 90:10 (Tables 1, 2). Although no differences in the AS:SS ratio were found between animals, comparison among layers revealed a statistically significant difference in layer 1 (χ^2^; P <0.0001), which displayed a higher proportion of AS than the other layers (96% AS and 4% SS; Table 2; Supplementary Table 1; Fig. 4B).

### Synaptic size

The study of the synaptic size was carried out analyzing the area of the SAS of each 3D reconstructed synapse (n=7,239) in the FIB/SEM stacks (Fig. 3C). To characterize the distribution of SAS area data, we performed goodness-of-fit tests to find the theoretical probability density functions that best fitted the empirical distributions of SAS areas in each layer and in all layers pooled together. We found that the best fit corresponded to log-normal distributions (Supplementary Fig. 2). These log-normal distributions, with some variations in the location (μ) and scale (σ) parameters

(Supplementary Table 2), were found in all layers for both AS and SS, although the fit was better for AS than for SS, probably due to the smaller number of SS.

The analysis of the SAS areas showed that AS were significantly larger than SS considering all layers (MW; p=0.0087; Fig. 4C, Tables 1, 2; Supplementary Table 1). These differences were also found in the frequency distribution analyses (KS; p<0.0001), showing that the proportion of small SAS areas were higher in SS than in AS (Fig. 4D). Analysis of the SAS area per layer showed that SAS areas of AS are larger than those from SS in all layers except in layer VI, where AS had smaller values than SS (MW, p<0.05, Table 2).

### Three-dimensional spatial synaptic distribution

To analyze the spatial distribution of the synapses, the actual position of each of the synapses in each stack of images was compared with a random spatial distribution model (Complete Spatial Randomness, CSR). For this, the functions G, K and F were calculated in the 18 stacks (Supplementary Fig. 3). We found that in half of the stacks (9 out 18) the spatial distribution of synapses was compatible with a random distribution. In the other half of the samples, a slight tendency for a regular pattern was detected by the G function, which identified slightly larger distances to the nearest neighbor than those expected by chance (Supplementary Fig. 3).

The mean distance from each synapse centroid to its nearest neighboring synapse within the counting frame was also calculated. Synapses that were closer to the boundaries of the counting frame than to any other synapse were excluded from the calculations, since their nearest neighbor could be placed outside the counting frame at an unknown distance (Baddeley et al., 1993; Illian et al., 2007). The estimated inter-synaptic distance was 591±33 nm (mean±SD) for all animals and layers. These measurements were calculated separately per layer, yielding the highest value in layer VI (631±43nm) and the lowest in layer I (532±68 nm; Table 2; Supplementary Table 1), although the differences were not statistically significant (KS, p<0.05).

## DISCUSSION

The present study constitutes the first description of the ultrastructural synaptic characteristics of the neuropil from the cerebral cortex of the Etruscan shrew. The following major results were obtained: (i) cortical synaptic density was very high, particularly in layer I; (ii) the vast majority of synapses were excitatory — the highest proportion was found in layer I; (iii) excitatory synapses were larger than inhibitory synapses in all layers except in layer VI; and (iv) synapses were either randomly distributed in space or showed a slight tendency for a regular pattern What follows is a discussion of the above results in comparison with data obtained from the human cerebral cortex (unless otherwise specified). From an evolutionary point of view, it is of particular interest to compare the synaptic organization of the brain of the smallest mammal with that of the much larger human brain, whose synaptic organization is thought to have reached the highest level of complexity. Fortunately, data is available from the human cerebral cortex that was obtained using the same methodology (Domínguez-Álvaro et al., 2018; 2021a; Cano-Astorga et al., 2021) as that used in the present study, avoiding the difficulties that are inherent when comparing different studies using different approaches. Thus, similarities and differences in the synaptic organization can be directly compared to examine what characteristics are conserved in evolution.

### Number of synapses and spatial distribution

Synaptic density is a useful parameter for describing synaptic organization, in terms of connectivity and functionality. In the Etruscan shrew, high densities of synapses were found in all layers of the somatosensory cortex, with a mean synaptic density of 1.31 synapses/μm^3^. No quantitative analysis of the synapses in the Etruscan shrew cerebral cortex has been performed previously and, thus, it is not possible to compare our results with those of others. However, the values for the synaptic density of the Etruscan shrew are almost triple those obtained in cortical samples from human temporal and entorhinal cortex using the same 3D EM method and image analysis (Cano-Astorga et al., 2021; Domínguez-Álvaro et al., 2021; Table 3).

**Table 3.**
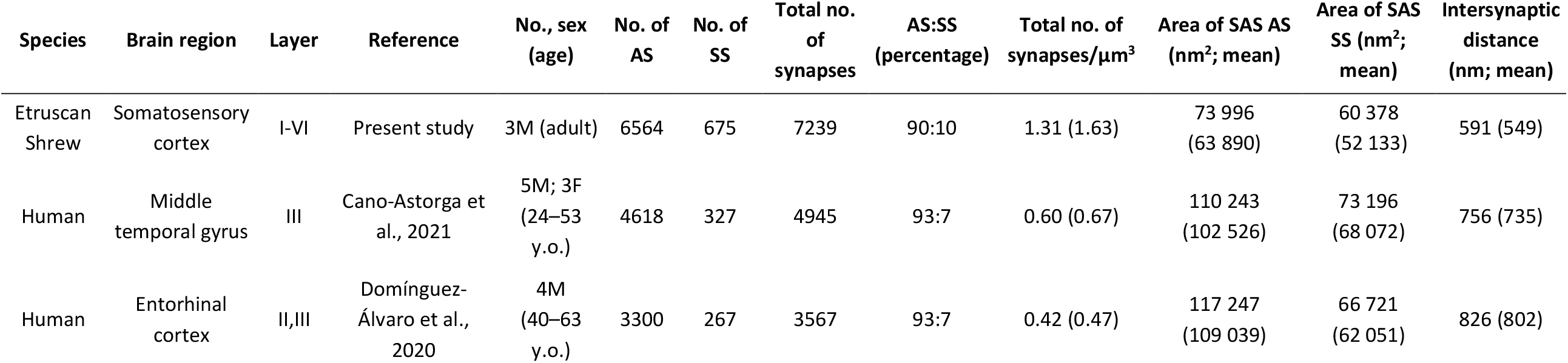
Summary of synaptic data from FIB/SEM studies for comparison. Data in parentheses are not corrected for tissue shrinkage. AS: asymmetric synapses; F: female; M: male; SAS: synaptic apposition surface; SS: symmetric synapses.

| Species | Brain region | Layer | Reference | No., sex (age) | No. of AS | No. of SS | Total no. of synapses | AS:SS (percentage) | Total no. of synapses/$\mu\text{m}^3$ | Area of SAS AS (nm<sup>2</sup>; mean) | Area of SAS SS (nm<sup>2</sup>; mean) | Intersynaptic distance (nm; mean) |
| --- | --- | --- | --- | --- | --- | --- | --- | --- | --- | --- | --- | --- |
| Etruscan Shrew | Somatosensory cortex | I-VI | Present study | 3M (adult) | 6564 | 675 | 7239 | 90:10 | 1.31 (1.63) | 73 996 (63 890) | 60 378 (52 133) | 591 (549) |
| Human | Middle temporal gyrus | III | Cano-Astorga et al., 2021 | 5M; 3F (24–53 y.o.) | 4618 | 327 | 4945 | 93:7 | 0.60 (0.67) | 110 243 (102 526) | 73 196 (68 072) | 756 (735) |
| Human | Entorhinal cortex | II,III | Domínguez-Álvaro et al., 2020 | 4M (40–63 y.o.) | 3300 | 267 | 3567 | 93:7 | 0.42 (0.47) | 117 247 (109 039) | 66 721 (62 051) | 826 (802) |

The highest synapse density was found in layer I (1.70 synapses/μm^3^; Table 2), which has a very low density of neurons (Fig. 1). In addition, the thickness of layer I in the Etruscan shrew somatosensory cortex represents about 20% of the total cortical thickness (Naumann et al., 2012). That is, in the Etruscan shrew, given the high synaptic density in layer I and its relatively large proportion, this layer greatly contributes to the total number of synapses in the somatosensory cortex.

In the present study, the AS:SS ratio was 90:10 (ranging from 88:12 to 96:4), which is within the range of the cortical values reported from other species. The percentage of AS and SS varies between 80–95% and 20–5%, respectively — in all the cortical layers, cortical areas and species examined so far using transmission electron microscopy (Beaulieu and Colonnier, 1985; Megías et al., 2001; DeFelipe et al., 2002; Bourne & Harris, 2011; DeFelipe, 2011, 2015) or FIB/SEM (Santuy et al., 2018a; Domínguez-Álvaro et al., 2018, 2021; Montero-Crespo et al., 2020; Cano-Astorga et al., 2021). However, layer I of the Etruscan shrew displays the highest proportion of AS (approximately 96:4, AS:SS) compared to other cortical layers where this proportion was similar (89:11). This suggests that there is a layer-specific excitatory-inhibitory balance.

Regarding the spatial organization of synapses, we found that the synapses either fitted to a random distribution in the neuropil or showed a slight tendency for a regular pattern, where points tend to separate from each other more than expected by chance. In the latter case, this may be because the spatial statistical functions are applied to the centers of gravity or centroids of the synaptic junction. However, it is important to take into account that synaptic junctions cannot overlap, and thus the minimum distances between their centroids are limited by the sizes of the synaptic junctions themselves, resulting in a slightly dispersed distribution of the centroids. This type of spatial distribution, which is based on a random distribution with a minimum-spacing rule, has also been found in the rat somatosensory cortex (Merchan-Perez et al., 2014; Anton-Sanchez et al., 2014) and several regions of the human brain including frontal cortex, transentorhinal cortex, entorhinal cortex, temporal cortex and CA1 hippocampal field (Blazquez-Llorca et al., 2013; Domínguez-Álvaro et al., 2018; 2021a; Montero-Crespo et al., 2020; Cano-Astorga et al., 2021). Therefore, the present results indicating the random spatial distribution of synapses are in line with the proposed widespread ‘rules’ of the synaptic organization of the mammalian cerebral cortex.

### Synaptic size

It has been proposed that synaptic size is directly related to neurotransmitter release probability, synaptic strength, efficacy and plasticity (e.g., Nusser et al., 1998; Ganeshina et al., 2004a; Tarusawa et al., 2009; Matz et al., 2010; Holderith et al., 2012; Südhof, 2012; Montes et al., 2015). Hence, the analysis of the synaptic size provides useful information about the synaptic function of a particular brain region.

In the present study, we used the values obtained from the SAS, which is equivalent to the interface between the active zone and the postsynaptic density (Morales et al., 2013). Thus, investigating SAS area is as an appropriate approach to analyze the synaptic size (Morales et al., 2013). Analysis of the somatosensory cortex of the Etruscan shrew has shown that SAS area was larger in AS than in SS (Fig. 4), which is similar to previous data obtained in other cortical areas and species using the same method (Domínguez-Álvaro et al., 2021; Montero-Crespo et al., 2020; Cano-Astorga et al., 2021). In addition, the SAS area of both types of synapses (asymmetric and symmetric) follows log-normal distributions, as do many other neuroanatomical and physiological variables such as synaptic strength, axonal width, and cortico-cortical connection density (Buzsaki & Mizuseki, 2014; Markov et al., 2014; Robinson et al., 2021).

However, we observed that the SAS area for AS was much smaller (73,996 nm^2^) compared to that found in the human temporal cortex and entorhinal cortex (110,243 nm^2^ and 117,247 nm^2^, respectively; Table 3). However, the SAS area for SS was similar to that found in other species and cortical regions (Table 3), which may indicate that SS are more homogeneous across species than AS (Santuy et al., 2018b).

It should be kept in mind that the SAS area of the AS is rather variable (Table 3). Larger and more complex synapses have been proposed to have more receptors in their postsynaptic elements than small synapses, and are thought to constitute a synaptic population with long-lasting memory-related functionality (e.g., Geinisman et al., 1993; Lüscher et al., 2000; Toni et al., 2001; Ganeshina et al., 2004a, 2004b) — whereas, small active zones may play a special role in synaptic plasticity (Kharazia & Weinberg, 1999). Thus, the presence of relatively small AS in the neuropil of the somatosensory cortex of the Etruscan shrew may indicate a lower release probability, synaptic strength and efficacy. In fact, hippocampal mossy fibers in Etruscan shrew have shown lower long- and short-term plasticity, as well as reduced expression of synaptotagmin-7 (a key synaptic protein in the regulation of presynaptic function) compared to mice (Beed et al., 2020). In this regard, it has been shown that mammalian brain synapses contain thousands of synaptic proteins resulting a high level of synapse diversity (Biederer et al., 2017; Zhu et al., 2018), which may result in synaptic species-specific differences (Curran et al., 2021). Thus, it is likely that molecular characterization of the synaptic proteins in the Etruscan shrew cortex may reveal additional specific synaptic characteristics.

### Layer-specific differences

In general, the structure of cortical layer 1 is highly conserved across cortical areas and mammalian species and it shows distinctive characteristics. It has sparse neurons, which are GABAergic interneurons (Schuman et al., 2019), and most of its volume is occupied by neuropil (Santuy et al., 2018c; Alonso-Nanclares et al., 2008). Layer 1 is the predominant input layer for top-down information, relayed by abundant projections that provide signals to the tuft branches of the pyramidal neurons (reviewed in Schuman et al., 2021). In particular, layer 1 receives axons from the thalamus and other cortical areas (cortico-cortical connections), as well as from local neurons from deeper layers (Muralidhar et al., 2014; Schuman et al., 2021). It has been proposed that layer 1 mediates the integration of contextual and cross-modal information in top-down signals with the input specific to a given area, enabling flexible and state-dependent processing of feedforward sensory input arriving deeper in the cortical column (reviewed in Schuman et al., 2021). In addition, layer 6 also showed some particular characteristics, including the lowest synaptic density and a lower SAS area for AS than SS. Thus, synaptic characteristics show layer-specific differences. However, the specific functional significance of the laminar differences in the synaptic organization of the Etruscan shrew remains to be elucidated.

In summary, certain general synaptic characteristics of the cerebral cortex of the Etruscan shrew are remarkably similar to those found in the human cerebral cortex including the following: (i) the vast majority of synapses are excitatory, (ii) the size of synaptic junctions follows a lognormal distribution, (iii) excitatory synapses are larger than inhibitory synapses and (iv) synapses fit quite closely to a random spatial distribution. Therefore, these synaptic characteristics might be considered as basic bricks of the cortical synaptic organization in mammals. Regarding the density and number of synapses, the Etruscan shrew has a high synaptic density of around 1,300 ×10^6^ synapses per mm^3^, which is almost triple the estimated synaptic density (about 500 ×10^6^ synapses per mm^3^) in the human cortex. Since the estimated volume of the Etruscan shrew cerebral cortex is 10.6 mm^3^ (Nauman et al., 2012), the total number of synapses would be about 14,000 ×10^6^, whereas in the human cortex this number can be up to 138,000,000 ×10^6^ synapses (based on a total cortical volume of 553,000 mm^3^, as reported by Ribeiro et al., 2013). That is, the cortical volume of the human brain is about 50,000 times larger than the cortical volume of the Etruscan shrew, but the total number of cortical synapses in human is ‘only’ around 20,000 times the number of synapses in the shrew. Furthermore, the synaptic junctions are about 35% smaller in the Etruscan shrew, which may be considered a relatively small difference. Thus, these differences in the number and size of synapses cannot be attributed to a brain size scaling effect, but rather to adaptations of synaptic circuits to particular functions.

## ACKNOWLEDGEMENTS

This work was supported by the following Grants: PGC2018-094307-B-I00 funded by MCIN/AEI/10.13039/501100011033; Cajal Blue Brain Interdisciplinary Platform (CSIC, Spain); and Research Fellowships funded by MCIN/AEI/10.13039/501100011033 for N.C.-A (PRE2019-089228) and S.P.-A. (FPU19/00007). We would like to thank Nick Guthrie for his helpful comments and excellent editorial assistance.

## AVAILABILITY OF DATA AND MATERIALS

Most data generated or analyzed during this study are included in the main text, the tables and the supplementary tables. Datasets used during the current study are available from the corresponding author on reasonable request.

## Abbreviation list

3D: three-dimensional
AS: asymmetric synapses
CF: counting frame
CSR: Complete Spatial Randomness
FIB/SEM: focused ion beam/scanning electron microscopy
KS: Kolmogorov-Smirnov
MW: Mann-Whitney
PB: phosphate buffer
PSD: postsynaptic density
SAS: synaptic apposition surface
SD: standard deviation
SEM: standard error of the mean
TEM: transmission electron microscopy
SS: symmetric synapses

## SUPPLEMENTARY MATERIAL

**Supplementary Figure 1.**
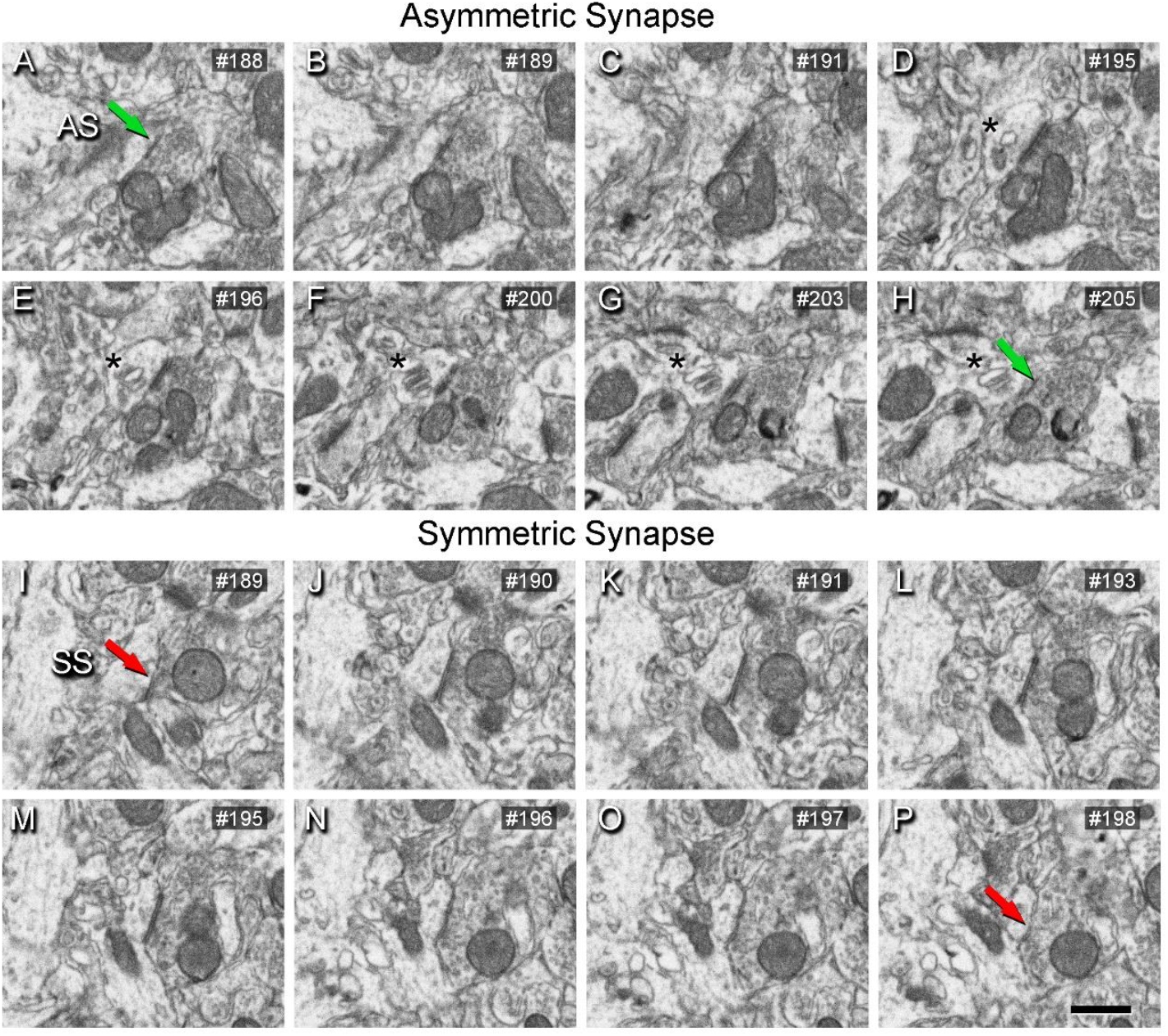
Sequence of FIB/SEM serial images of an AS (A–H) and an SS (I–P) indicated in Figure 2. Numbers on the top right of each panel indicate the number of each section from a stack of serial sections. Synapse classification was based on the examination of full sequences of serial images, see “Three-dimensional analyses of synapses” for further details. Asterisks (in D–H) indicate a spine apparatus in a postsynaptic dendritic spine head. Scale bar shown in P represents 500 nm in A–P.

**Supplementary Figure 2.**
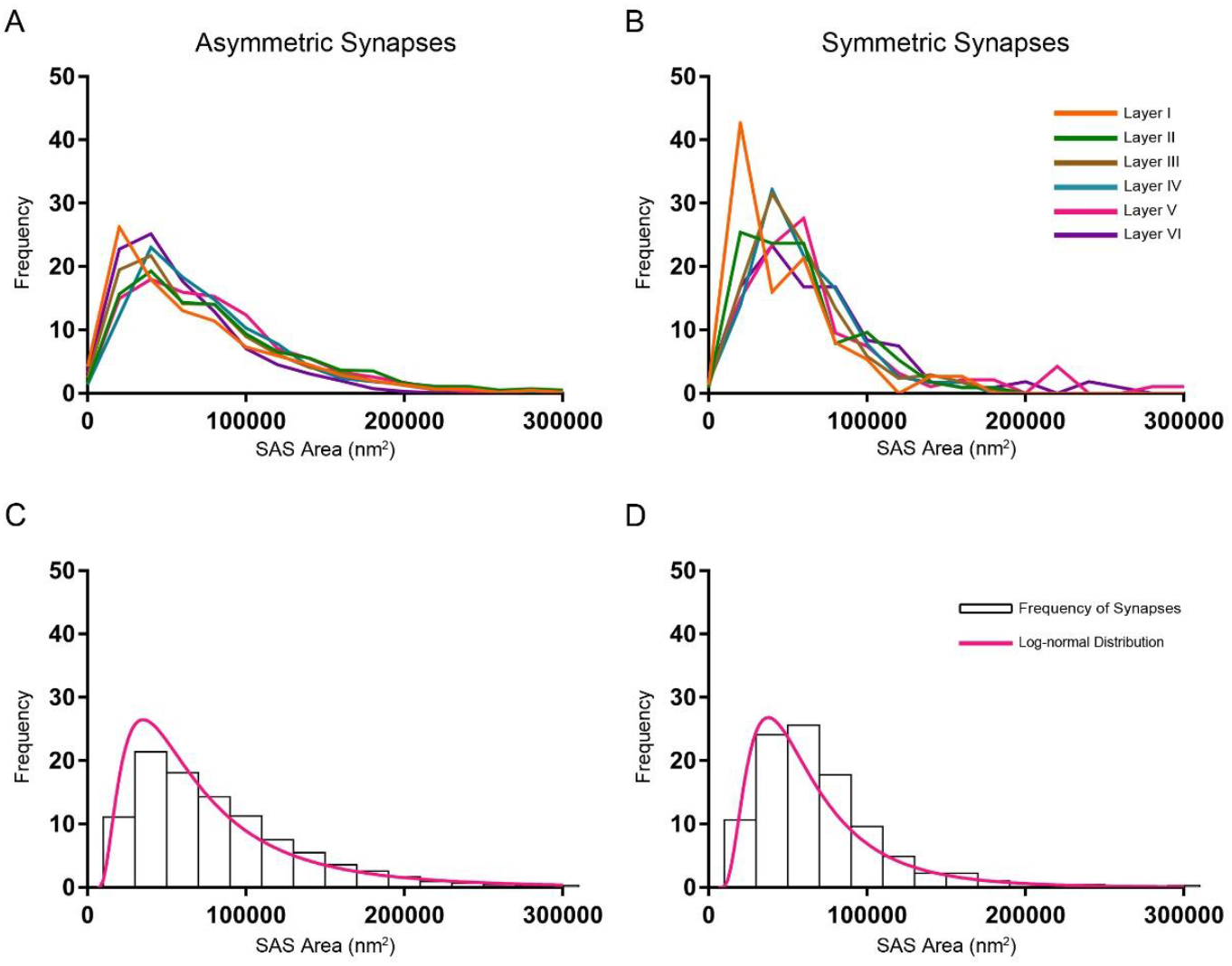
Frequency histograms of SAS areas and their corresponding best-fit probability density functions. (**A, B**) Frequency histograms of SAS areas in the six cortical layers are represented for AS and SS in A and B, respectively. (**C, D**) Frequency histograms (white bars) and best-fit distributions of the theoretical probability synaptic density functions (magenta traces) have been represented. The best-fit probability functions were log-normal distributions. Curve fitting was always better for AS (C) than for SS (D), probably because of the smaller sample size of SS (Supplementary Table 2). The parameters μ and σ of the log-normal curves are shown in Supplementary Table 2.

**Supplementary Figure 3.**
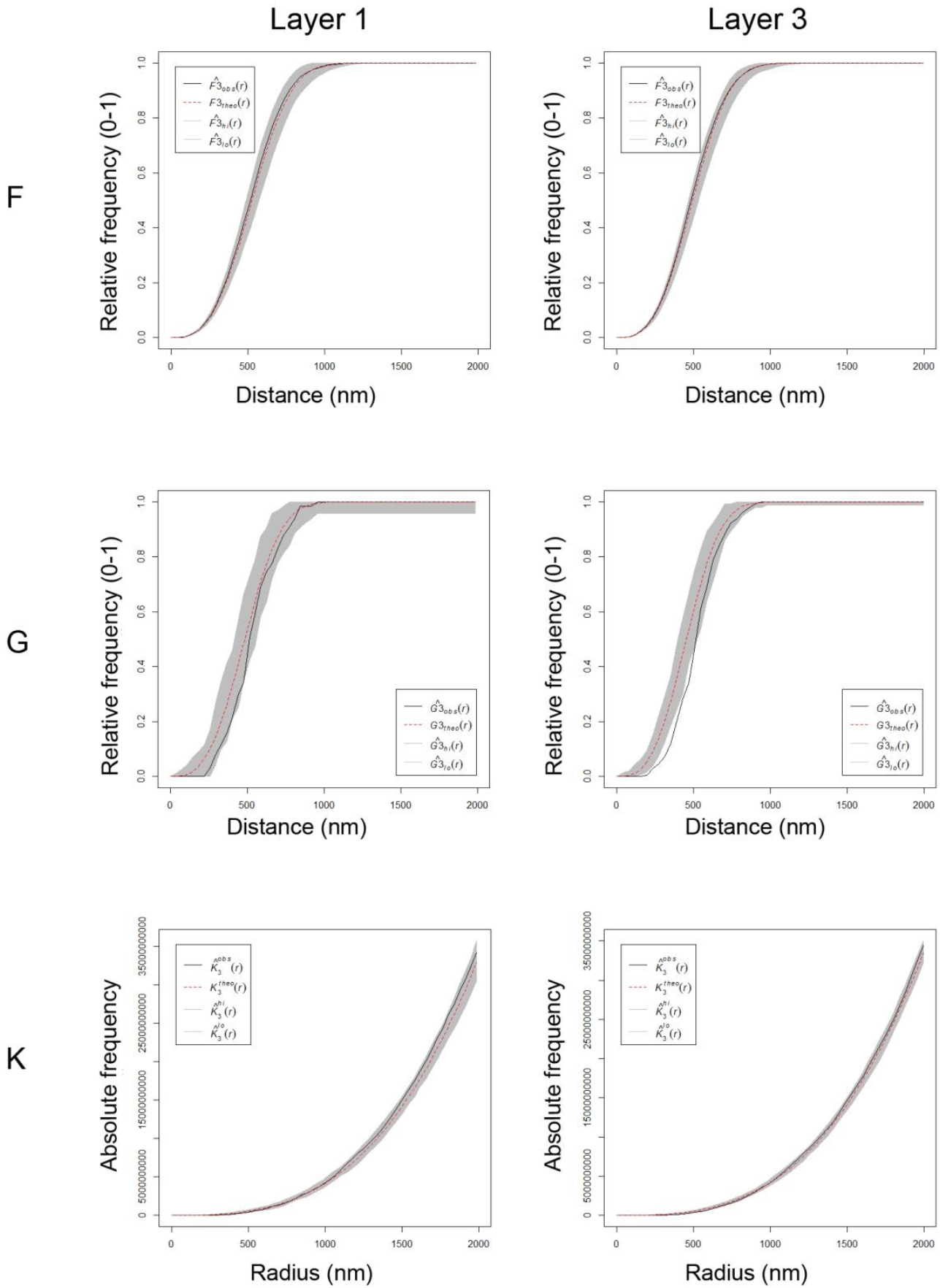
Analysis of the 3D synaptic spatial distribution in somatosensory cortex from the Etruscan shrew. Red dashed traces correspond to a theoretical homogeneous Poisson process for each function (F, G, K). The black continuous traces correspond to the experimentally observed function in the sample. The shaded areas represent the envelopes of values calculated from a set of 99 simulations. Plots show a distribution which fits into a Poisson function, but the experimental function from layer 3 for the G-function is partially out of the envelope. Plots obtained in layer 1 and layer 3 from animal MS1.

**Supplementary Table 1.** Data per layer and animal. Data in parentheses are not corrected for shrinkage. AS: asymmetric synapses; SAS: synaptic apposition surface; SEM: standard error of the mean; SS: symmetric synapses.

| Layer | Animal | No. of AS | No. of SS | Total no. of synapses | CF Volume ($\mu\text{m}^3$) | No. of AS /$\mu\text{m}^3$ | No. of SS /$\mu\text{m}^3$ | No. of synapses /$\mu\text{m}^3$ | Area of the SAS of AS ($\text{nm}^2$; mean ± SEM) | Area of the SAS of SS ($\text{nm}^2$; mean ± SEM) | Distance to the nearest neighbor (nm; mean) |
| --- | --- | --- | --- | --- | --- | --- | --- | --- | --- | --- | --- |
| I | MS1 | 267 | 10 | 277 | 233 (187) | 1.15 (1.43) | 0.04 (0.05) | 1.19 (1.48) | 92532±5536 (79895±4780) | 56229±17270 (48550±14912) | 607 (564) |
| | MS2 | 346 | 56 | 402 | 222 (178) | 1.99 (2.47) | 0.08 (0.10) | 2.06 (2.57) | 64930±2489 (56062±2149) | 43780±6619 (37801±5715) | 474 (440) |
| | MS3 | 446 | 55 | 501 | 434 (348) | 1.74 (2.17) | 0.11 (0.14) | 1.85 (2.31) | 64560±2090 (55743±1804) | 46323±4737 (39997±4090) | 517 (480) |
| II | MS1 | 250 | 58 | 308 | 330 (265) | 1.05 (1.31) | 0.17 (0.21) | 1.22 (1.52) | 77887±3077 (67250±2657) | 58300±3510 (50338±4411) | 602 (559) |
| | MS2 | 184 | 36 | 220 | 262 (210) | 1.25 (1.56) | 0.14 (0.17) | 1.39 (1.73) | 83599±3510 (72182±3031) | 53328±3637 (46045±4588) | 569 (529) |
| | MS3 | 257 | 34 | 291 | 218 (175) | 1.45 (1.80) | 0.10 (0.13) | 1.55 (1.93) | 89448±3637 (77232±3140) | 58128±5873 (50190±5071) | 554 (515) |
| III | MS1 | 441 | 17 | 458 | 362 (291) | 1.23 (1.53) | 0.15 (0.19) | 1.38 (1.72) | 81420±2811 (70301±2427) | 61116±2468 (52769±4457) | 569 (529) |
| | MS2 | 327 | 36 | 363 | 405 (325) | 1.03 (1.29) | 0.15 (0.18) | 1.18 (1.47) | 72512±2468 (62609±2131) | 49283±1998 (42553±3061) | 599 (556) |
| | MS3 | 418 | 60 | 478 | 336 (269) | 1.64 (2.04) | 0.17 (0.21) | 1.81 (2.25) | 63716±1998 (55014±1725) | 62635±4534 (54081±3914) | 523 (486) |
| IV | MS1 | 283 | 24 | 307 | 353 (283) | 0.71 (0.88) | 0.16 (0.20) | 0.87 (1.09) | 70369±3131 (60758±2704) | 56759±2811 (49008±3332) | 621 (577) |
|  | MS2 | 391 | 30 | 421 | 295 (237) | 0.96<br>(1.19) | 0.08<br>(0.10) | 1.04<br>(1.30) | 79449±2811<br>(68599±2427) | 55917±2736<br>(48280±5886) | 635 (590) |
|  | MS3 | 363 | 42 | 405 | 217 (174) | 1.56<br>(1.94) | 0.15<br>(0.18) | 1.71<br>(2.13) | 76698±2736<br>(66224±2362) | 57249±4820<br>(49431±4161) | 568 (528) |
| V | MS1 | 754 | 48 | 802 | 251 (201) | 0.73<br>(0.91) | 0.14<br>(0.18) | 0.88<br>(1.09) | 87083±4144<br>(75190±3578) | 80928±2258<br>(69876±9054) | 711 (661) |
|  | MS2 | 316 | 22 | 338 | 377 (302) | 1.04<br>(1.29) | 0.08<br>(0.10) | 1.12<br>(1.39) | 67506±2258<br>(58287±1950) | 53011±2612<br>(45771±6598) | 585 (544) |
|  | MS3 | 550 | 56 | 606 | 284 (228) | 1.22<br>(1.52) | 0.10<br>(0.12) | 1.32<br>(1.64) | 83137±2612<br>(71783±2255) | 79508±11278<br>(68649±9737) | 609 (566) |
| VI | MS1 | 339 | 32 | 371 | 301 (241) | 0.85<br>(1.06) | 0.11<br>(0.14) | 0.97<br>(1.21) | 54205±2528<br>(46802±2183) | 69309±1973<br>(59843±7435) | 664 (617) |
|  | MS2 | 346 | 28 | 374 | 395 (317) | 0.92<br>(1.14) | 0.11<br>(0.13) | 1.02<br>(1.28) | 56034±1973<br>(48381±1704) | 64423±2453<br>(55625±6337) | 583 (542) |
|  | MS3 | 286 | 31 | 317 | 301 (242) | 0.95<br>(1.18) | 0.10<br>(0.13) | 1.05<br>(1.31) | 66843±2453<br>(57715±2118) | 80585±9259<br>(69580±7995) | 647 (601) |

**Supplementary Table 2.** Number of synaptic SAS analyzed (n), the location (μ) and scale (σ) of the best-fit log-normal distributions in the six cortical layers. AS: asymmetric synapses; SAS: synaptic apposition surface; SS: symmetric synapses.

|  | AS |  |  | SS |  |  |
| --- | --- | --- | --- | --- | --- | --- |
| | *n* | $\mu$ | $\sigma$ | *n* | $\mu$ | $\sigma$ |
| Layer I | 1462 | 10.79 | 0.90 | 75 | 10.50 | 0.73 |
| Layer II | 989 | 11.05 | 0.80 | 114 | 10.75 | 0.65 |
| Layer III | 1414 | 10.91 | 0.79 | 171 | 10.80 | 0.59 |
| Layer IV | 872 | 11.03 | 0.68 | 114 | 10.81 | 0.56 |
| Layer V | 921 | 11.02 | 0.73 | 94 | 10.93 | 0.71 |
| Layer VI | 906 | 10.74 | 0.74 | 107 | 10.93 | 0.72 |
| Layers I–VI | 6564 | 10.91 | 0.80 | 675 | 10.80 | 0.66 |

